# Lane-maze for preference testing in flies

**DOI:** 10.1101/2021.07.18.452790

**Authors:** Fabiola Boz Eckert, Dhiozer de Brittos Valdati, José Marino Neto, Daniela Cristina de Toni, Cilene Lino de Oliveira

**Affiliations:** UFSC - Postgraduate Program in Pharmacology (Florianópolis, SC-Brazil); UFSC - Institute of Biomedical Engineering (Florianópolis, SC-Brazil); UFSC - Department of Cellular Biology, Embryology and Genetics (Florianópolis, SC-Brazil); UFSC - Department of Physiological Sciences (Florianópolis, SC-Brazil)

**Keywords:** behaviour, motor activity, preference, replacement, sexual differences

## Abstract

*Drosophila melanogaster* is a candidate species to replace rodents in some neurobiological studies, encouraging attempts to develop behavioural tests for these flies. This study aimed to develop a behavioural test to simultaneously evaluate ethological (categorical) aspects of the motor and fluid intake activities, which may be used to assess sucrose preference in flies. For that, a lane-maze was 3D-printed to accommodate up to 14 individual flies in a single trial. Each lane had a capillary filled with 5% sucrose solution attached to one of the extremities. To validate a 5-min lane-maze test, male and female flies (adults, 5-6 days of age) underwent 0, 2, 8 or 20 h of food deprivation (FD, n=9-11/group) before testing. Duration of locomotion, immobility and grooming in the lane or capillary were scored from the video-recorded trials using EthoWatcher software. Minor effects of sex or FD were observed in the behaviours of flies. Independent of sex or FD, flies spent proportionally longer on the capillary than on the lane. Flies exhibited a significantly higher preference than expected for the capillary zone when food-deprived for 2h (males) or 20 h (females). Data suggest that short lane-maze test is a feasibly high throughput assessment of sucrose preference in flies, which may be sexually dimorphic as in other species studied so far.

## Introduction

Models and behavioural tests in rodents have been criticised by the scientific community due to concerns about their clinical validity, application, and animal welfare, ^1^ pushing the research field to find alternatives to the use of vertebrates in research. The 3Rs principles may help the scientific community to perform better animal research.^2^ Refinement may minimise animal suffering and improve welfare, while reduction may guide methods minimising the number of animals used per experiment. The principle of replacement can lead to methods with full or partial substitution of animals using alternative methods. Full replacement avoids using any non-human animal in research, by substituting them with human volunteers, tissues and cells, mathematical and computer models, and established cell lines. Partial replacement avoids using vertebrates in research, by substituting them with invertebrates such as nematode, social amoebae, flies, and others, which cannot experience suffering based on current scientific knowledge although they react to threats.

In this context, *Drosophila melanogaster* has been considered a candidate organism to replace laboratory rodents or other vertebrates in neuroscientific studies.^3,4,5,6^ Feeding, sleep, aggression, preferences, learning, memory and responses to stress in *Drosophila* are controlled by monoamines, distinct monoamine receptors and neural circuits.^5,6, 7,8,9^ Beyond that, *D. melanogaster* has well-known anatomy, physiology, genome, proteome and natural behaviour.^3,10^ Notably, different strains of *Drosophila* with various phenotypes may be kept and reproduced quickly in the laboratory allowing for the planning of well-powered experiments as required in neurobiological research. Nevertheless, unlike rodents, models, behavioural tests for studying psychopharmacology or neurobiology of stress in *Drosophila* are still in the embryonic steps of standardisation. ^11,12^ In laboratory rodents, symptoms of stress may be assessed using several behavioural tests such as positive or negative avoidance, forced swimming, tail suspension, sucrose preference and others.^13,14,15,16, 17^

Since the pioneer study reporting learned helplessness in flies,^18^ several attempts to develop models and behavioural tests for stress response in *Drosophila* have been made. ^19, 20^ Flies trained to receive inescapable heat shocks failed to escape from the stressor even when the opportunity appeared, different from those trained with escapable heat shocks.^18^ Another study demonstrated that exposure to vibrations for three days reduced the walking of flies in the climbing test.^5^ A random sequence of variable stresses (including cooling, heating, sleep and food deprivation) for ten days induced high immobility in the forced swimming test, aggressiveness in social interaction test and anhedonia-like behaviour in a sucrose preference test in *Drosophila*.^19^ A protocol using a short, random and unpredictable combination of stressors (18 h-social isolation, six hour-fasting, 20 min-heat shock and five min-electric shocks) induced hyperactivity and *centrophilia* (preference for staying in the centre of the arena) in an open field test, as well as less preference for sucrose in a one-minute preference test in a Petri dish.^20^ Fasting *per se* induced hyperactivity and centrophilic behaviour in flies but, in contrast to the combination of stressors, increased sucrose preference ^20^, probably, due to a nutritional deficit. Although signs of stress in *Drosophila* seemed to be stressor-dependent, different types of stimuli affected motor activity in the open field test and the preference for sucrose.

Preference for sucrose may be assessed using tests measuring sucrose intake in *Drosophila*, such as fly Proboscis and Activity Detector (flyPAD) ^21^, capillary feeding (CAFE),^22, 23^ proboscis’s extension,^24^ dyed abdomen,^25^ Fly Liquid-Food Interaction Counter (FLIC),^26^ or “Activity Recording Capillary Feeder” (ARC). ^27^ Except for ARC ^27^, most of the abovementioned methods were developed to investigate a single experimental unit at a time (an individual fly or a group of flies). ARC ^27^ employed positional tracking allowing for the simultaneous assessment of sleeping and fluid intake from up 60 individual flies in the same trial. Inspired by ARC ^27^, the aim here is to develop an apparatus allowing for the analysis of the ethological (categorical) aspects of motor activity and fluid intake in several experimental units simultaneously. Then, an apparatus with 14 parallel lanes (lane-maze) was printed in plastic to accommodate up to 14 individual flies in a single behavioural trial. ^28^ In both extremities of each lane, there is a hole to insert a capillary that may be filled with palatable fluid, which may be used to simulate sucrose preference tests as performed in rodents.

In the field of psychopharmacology, behavioural tests in rodents like forced swimming, tail suspension, and others are habitually short (5 to 15 minutes). ^13,14,15,16,17^ Therefore, here, a short version of a lane-maze test (5 minutes) was tried. To validate the 5-min lane-maze test, adult, virgin, male and female flies undergone 0, 2, 8 or 20 h of food deprivation before behavioural testing. An extremity of every lane had a capillary filled with 5% sucrose solution attached, simulating a single-bottle preference test for rodents.^16^ Food deprivation is expected to induce hyperactivity and increased sucrose preference in the flies ^20^, which may be accessed by scoring the categorical aspects of behaviour of flies in the lane or capillary zones. These approaches should allow for high throughput testing to plan well-powered studies to investigate the behavioural signs of stress in *D. melanogaster*.

## Materials and Methods

### Flies

*Drosophila melanogaster* (Canton-S) used in this study were obtained from Stock Center Tucson (Arizona, USA). Both sexes were kept in a vivarium with controlled temperature and photoperiod (20 ± 1 °C, 12 h light-dark cycle, lights on at 6 am, 60-80% of relative humidity) in the Biomedical engineering laboratory at the Federal University of Santa Catarina. The flies were kept in “house-glasses” (300 mL glass bottles) containing standard food (supplementary methods) sealed with a foam stopper until hatching.

### Experimental design and procedures

On the day of eclosion from the pupa, flies were collected, anaesthetised on ice (−4°C) for ±1 min and separated by sex into different small tubes (7.5 cm x 1.1 cm) where virgin male or female flies were maintained in groups of 8-10, under identical medium conditions (1 g/tube) for five days before allocation to the groups (control, C or food deprivation, FD). Flies were transferred from “house-glasses” to experimental tubes with food (C) or without food containing a paper filter saturated with water (FD). Experimental tubes were kept under vivarium conditions until the behavioural tests. For preliminary studies only flies of C groups were analysed. For experiments, flies of C or FD groups were kept in the experimental tubes for two (experiment 1), eight (experiment 2) or twenty hours (experiment 3) before behavioural testing. Each independent experiment was carried out on several experimental days to complete the final sample size. An experimental day consisted of testing up to twenty flies, ten males and ten females of C or FD groups simultaneously in the lane-maze. Afterwards, flies were euthanised by immersing the tubes in ice (−4°C, for ±5 min). See Figure 1B for the timeline.

**Legend for Figure 1:**
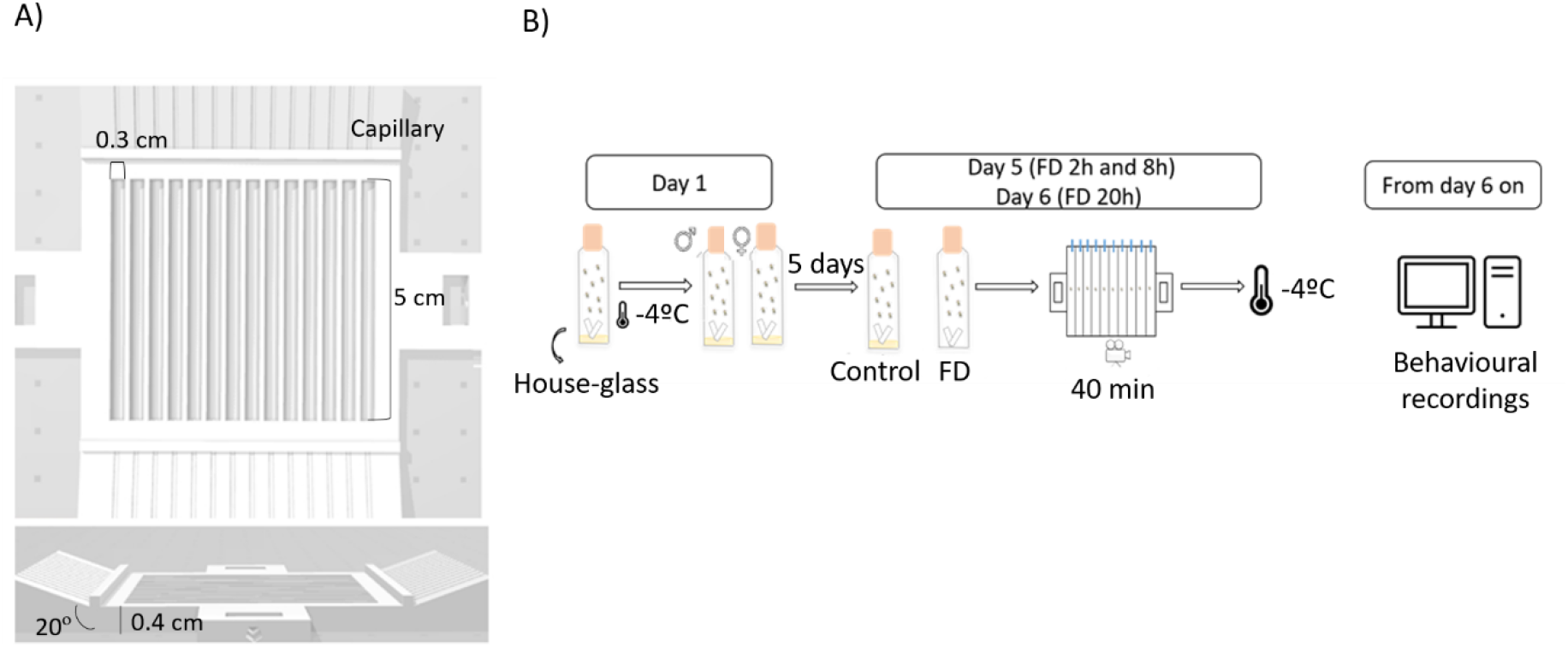
A-Lane-maze in a top view (upper figure) and a perspective view (lower figure); B-Experimental design. Abbreviations: FD= food deprivation, C=control.

#### Lane-maze test

Lane-maze ^28^ (Figure 1 A, supplementary methods) adapted from previously described methods ^22,23, 27^, consisted of a squared arena (external dimensions: 6.6 x 9.2 cm, internal dimensions: 5 x 5.5 cm) with 14 internal subdivisions (lanes) of identical dimensions (length: 5 cm, width: 0.3 cm) covered in a transparent acrylic plate. At both extremities of each lane, a hole (diameter: 0.05 cm) allowed for the insertion of a glass capillary at an angle of 20 degrees provided by the inclined edges of the external arena. Capillaries allow access to palatable fluids during behavioural tests carried out in a homemade test environment, consisting of a plastic bench adapted to support illumination (a string of LED lamps, 490 lux) and a camera (a USB Digital Microscope Camera, Lenovo ®) placed 42 cm above the lane-maze, allowing for simultaneous recording of 10 lanes. The lane-maze was maintained in a fixed position relative to the camera from test to test, on a platform (30 cm length, 4.5 height) made from white translucent acrylic with a pair of plastic sockets to fit the lane-maze. All was covered with an opaque screen to avoid illumination variations.

Procedures for testing in the lane-maze were as follows: 1-The lane-maze was placed on an ice-plate for ± 5 min. 2-Flies were anaesthetised on ice (−4°C) for ± 1 min and transferred carefully, using tweezers, to the respective lane of the cold lane-maze. 3-Each lane had a glass capillary (diameter: 0.04 cm) filled with blue sucrose solution (5%, dissolved in filtered water plus 0.1 mL of blue dye, food-grade colouring, Arcolor®, São Paulo-SP, Brazil) inserted in one of the extremities. Blue dye was added to the sucrose solution to facilitate the capillary visualisation in the video recordings and recognition after behavioural testing if flies had ingested sucrose. 4-The lane-maze was covered with the acrylic plate before flies recovered from the anaesthesia and were transferred to the platform of the test; 5-Video camera recording started at insertion of the lane-maze into the test environment. In the pilot study, behaviours were scored during the whole duration of the test (40 min-analysis), however, in the experiments 1, 2 and 3 behavioural testing began at five minutes after each fly moved for the first time in the lane (an indication that anaesthesia was over). The tests lasted a maximum of 40 min and were performed between 12 am and 4 pm in a room at 23±3 °C. 6-After the testing, flies were euthanised on ice (−4°C, ± 5 min). 7-After euthanasia, flies were examined to check for the presence of blue dye in their abdomen. 8-behavioural registering occurred after finishing the experiments.

### Data collection and analysis

Behavioural outcomes in the lane-maze were defined in the preliminary studies (see supplementary methods for detailed description). Behavioural registering was performed using Ethowatcher open-source software package^29^ available upon request. Raw data (duration (s), frequencies, latencies (s)) were analysed directly or used to calculate a “normalised duration” for every behavioural outcome registered on the lane or capillary (s/mm, proportion of occupancy) and a “preference index” (deviation from the “expected” occupancy). Normalisations (duration (s) divided by the area of the capillary or lane (mm)) were necessary to fit the duration of behaviour occurring in the region of the capillary (^~^5 mm) and lane (^~^45 mm) on the same scale. Thus, normalised duration of each behaviour provided a proportional occupancy of the capillary and other regions of the lane. Raw data and detailed calculations are provided in the supplementary results.

Raw behavioural outcomes, “normalised duration” or “preference indexes” were not normally distributed (significant Kolmogorov-Smirnov test) or presented homogeneous variances (significant Levene and Brown-Forsythe tests) even after different data transformations such as box-cox, logarithmic +1, logarithmic10 +1, square-root and square-root +0.5 (see supplementary results). Therefore, Kruskal–Wallis, a non-parametric one-way analysis of variance, was used to compare the four groups (F C, F FD, M C, M FD; female=F; male=M; control=C, food deprived= FD) in the independent experiments (Experiment 1, 2 or 3). *Post hoc* Mann-Whitney was used to assess significant differences between the two groups. For each independent experiment, the statistical analysis was performed blind for sex and treatment. Data are expressed as the mean(s)±standard error (SE). Comparisons with a p-value<.05 were considered statistically significant. These analyses were performed using the software Statistica (Statistica Inc, version 8).

“Preference indexes” for every independent experiment (Experiments 1, 2 or 3) were analysed by comparing expected and observed indexes of preference for the capillary region of the lane for every experimental group (F C, F FD, M C, M FD; female=F; male=M; control=C, food deprived= FD) using Sign test and Wilcoxon matched-pairs signed-ranks test.^30^ Data are expressed as the mean (%) ± standard error (SE). Comparisons with a p-value<.05 were considered statistically significant. These analyses were performed using the software Excel (Microsoft Office, version 16.37).

## Results

### Pilot studies: behavioural outcomes in the lane-maze

Preliminary analyses were performed using data of flies from control groups, see supplementary material for raw data, means, standard errors and standard deviations. Behavioural outcomes in the lane-maze were selected according to the quality and reliability of their assessment, which varied from almost perfect (Cohen’s *kappa* upper 80%), substantial (Cohen’s *kappa* upper 75%) to weak (Cohen’s *kappa* below 50%) (see supplementary material for detailed description). Only outcomes considered reliable (Cohen’s *kappa* upper 75%) were included in the quantitative analysis, as follows: immobility on the lane; locomotion on the lane; grooming on the lane; immobility on the capillary; locomotion on the capillary; grooming on the capillary. Time of behaviours were arbitrarily selected for further analysis because their baseline seemed less variable across experiments than latencies or frequencies.

Control flies spent more time in the lane than in the capillary region. Throughout a 40 min-test (Figure 2), the duration of locomotion (left panel), immobility (middle panel) and grooming (right panel) varied in flies of both sexes (females, upper panel; males, lower panel). After recovery from anaesthesia, i.e., after 5 min in the lane-maze, flies of both sexes displayed locomotion and grooming less prevalent than immobility (Figure 2). Considering that the flies seem to be under anaesthesia during the first 5 min in the lane-maze, the period before the 5^th^ min of the test was discarded from the analyses in the next experiments. After 20 min in the lane-maze, flies were mostly immobile while grooming decreased steadily until reaching stability at low levels. Locomotion seemed stable over the entire period of the test. Thus, the time window between the 5^th^ and 10^th^ minutes of the test was selected for the analysis in the next experiments.

**Legend for Figure 2:**
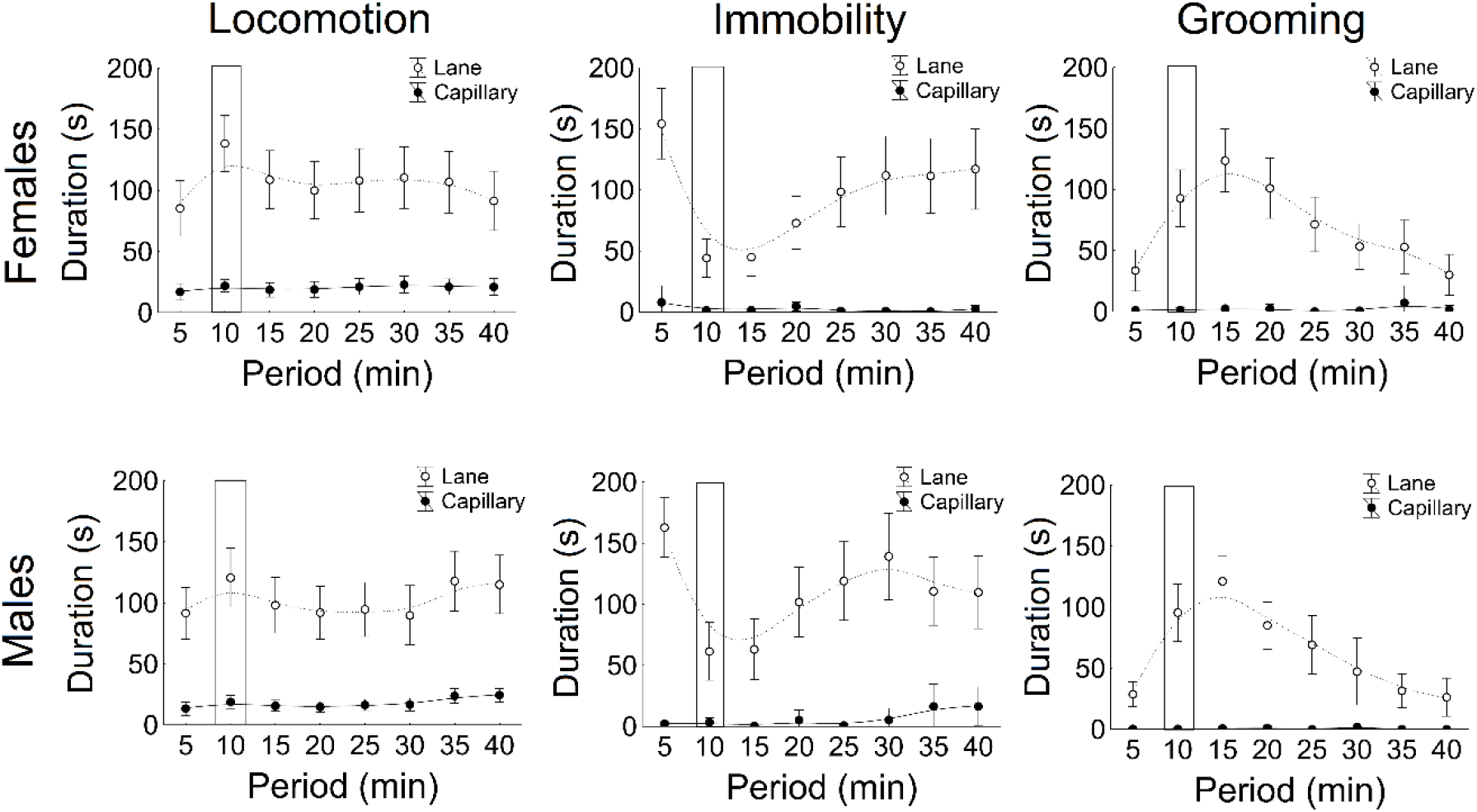
Duration (s) of locomotion (left panel), immobility (middle panel) and grooming (right panel) in the lane (white circles) or on the capillary (black circles) of females (upper panels, n=30) or males (lower panels, n=28) flies kept in standard laboratory conditions before the 40-min lane-maze test. Data are expressed as mean ± SEM. Each line represents a minimum square fit (distance-weighted minimum square fit) for the mean (Stiffness ± 25%). The black rectangles depict the period of the behavioural testing selected for data analysis in the next experiments (5^th^-10^th^ min).

### Effects of food deprivation on behaviour in the lane-maze

In experiments 1, 2 and 3, lane-maze test lasted 5-min starting after flies recovered from anaesthesia. For statistical results of non-significant comparisons of the experiments 1, 2 or 3, see supplementary results. In experiment 1 (2 h FD), no significant differences were observed among the groups (male or female flies of C or 2 h FD, Figure 3, upper panel) when analysing either raw or normalised duration of locomotion, immobility or grooming in the lane-maze test. In experiment 2 (8 h FD), except for normalised duration in the lane of immobility (H (3) =7.94 p =.04) and grooming (H (3) =8.5, p =.03), no significant differences were observed among the groups (male or female flies of C or 8 h FD, Figure 3, middle panel) when analysing either raw or normalised time of locomotion in the lane or capillary, immobility or grooming on the capillary. *Post hoc* Mann-Whitney indicate significant differences (p<.05) between the following comparisons: females of the C group had more grooming in the lane than FD; males of the FD group had more immobility in the lane than C. In experiment 3 (20 h FD), no significant differences were observed among the groups (male or female flies of C or 20 h FD, Figure 3, lower panel) when analysing either raw or normalised locomotion durations immobility or grooming in the lane-maze test.

**Legend for Figure 3:**
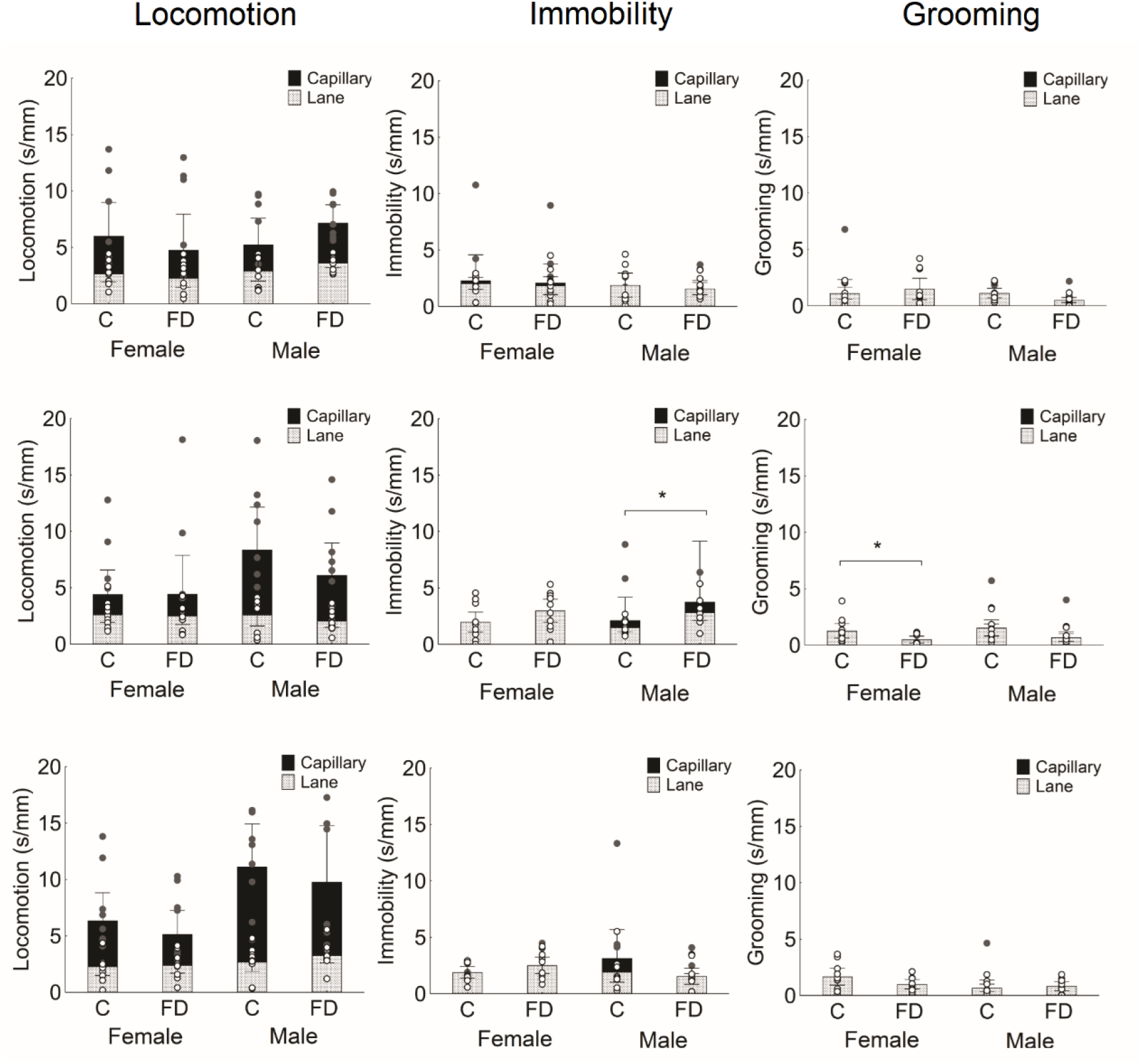
Proportion (seconds *per* millimetres, s/mm) of locomotion (left panels), immobility (middle panels), grooming (right panels) of female or male flies of the control (C) or food-deprived (FD) groups on the capillary (black bars) or lane (grey bars) during 5-min of the lane-maze test. Upper panel depicts result experiment 1 (C, females, n=9; males, n=9; 2 h FD, females, n=10; males, n=9); middle panel depicts result experiment 2 (C, females, n=11; males, n=9; 8 h FD, females, n=10; males, n=10); lower panel depicts result experiment 3 (C, females, n=10; males, n=10; 20 h FD, females, n=10; males, n=10). Data are expressed as mean ±SEM. Grey circles= individual data for the capillary, white circles= individual data for the lane. (*) significant according to the Mann-Whitney test, p<.05.

### Effects of food deprivation on capillary preference in the lane-maze test

Occupancy of the capillary region, i.e., “preference index”, varied from group to group depending on the sex of the flies and experimental conditions (Figure 4, table S6). In the C groups, the preference index of female flies was not significantly higher than expected in any experiment (Sign test, p>.05; Wilcoxon test, p>.05), while for males, it varied from experiment to experiment. In experiments 1 and 3, the preference index of male flies of C group was not significantly higher than expected (Sign test, p>.05; Wilcoxon test, p>.05). Although in experiment 2, the preference index of male flies of C group was significantly high (Sign test, p<.05; Wilcoxon test, p<.05). In FD groups, female flies had a significantly higher preference for the capillary region of the lane than expected when fasting lasted for 20 h (Experiment 3; Sign test, p>.05; Wilcoxon test, p<.05) but not for 2 or 8 h (Experiments 1 and 2; Sign test, p>.05; Wilcoxon test, p>.05). By contrast, male flies undergoing FD for 2 h (Experiment 1) had a significantly higher preference for the capillary region than expected (Sign test, p>.05; Wilcoxon test, p>.05). Longer times of FD (8 h or 20 h) increased preference for the capillary region, but not significantly (Sign test, p>.05; Wilcoxon test, p>.05).

**Legend for Figure 4:**
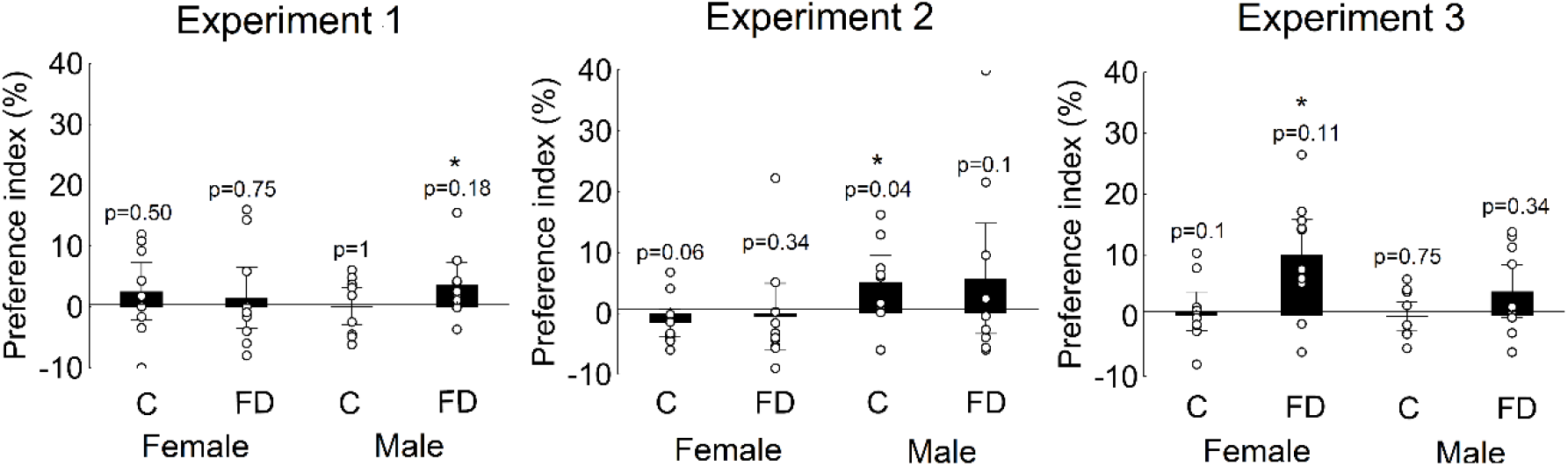
Preference for the capillary region of the lane (“preference index” %) of flies of control (C) or food-deprived (FD) groups in the experiment 1 (left panel; C, females, n=9; males, n=9; 2 h FD, females, n= 10; males, n=9), experiment 2 (middle panel, C, females, n=11; males, n=9; 8 h FD, females, n= 10; males, n=10) and experiment 3 (right panel, C, females, n=10; males, n=10; 20 h FD, females, n=10; males, n=10). Data are expressed as mean ±SEM. White circles= individual data, *p values* are of the Sign test, (*) significant according to Wilcoxon test, p<.05.

## Discussion

In this study, a lane-maze test was designed and standardised to simultaneously evaluate behaviours related to general motor activity and fluid intake in several individual flies. In flies, motor activity ^20, 31, 32, 33, 34^ and fluid intake ^21, 22, 23, 24, 25, 26^ are often measured independently in separate apparatuses, except ARC. ^27^ Motor activities are often measured using automatic tracking in the open field tests,^20, 31, 32, 33, 34^ while sucrose preference may be assessed in various analyses of fluid intake (e.g., CAFE ^22,23^; proboscis extension ^24^; ARC ^27^). Using positional tracking of individual flies and microcapillaries, ARC^27^ provides a high throughput platform for the simultaneous assessment of activity, immobility, and fluid intake. Similar to ARC,^27^ the lane-maze enabled the concurrent testing of multiple experimental units allowing for larger sample sizes, facilitating the design of well-powered studies. Indeed, the lane-maze enabled the evaluation of categorical aspects of activity like locomotion, immobility, grooming or fluid intake in flies in the same trial. Besides, lane-maze allocates a capillary in both sides of the lane. This is an advantage of this apparatus because allows the assessment of a binary preference measures, such as preference for different tastes or odors, in multiple experimental units simultaneously. Therefore, this apparatus could accelerate the acquisition of data in preference tests. The paired assessment of different outcomes in a single trial may reduce the time required for a behavioural experiment matched with individual genetic, proteomic, or biochemical profiles of individual animals.

Assessment of locomotion or immobility of flies was feasible and reliable in the different sectors of the lane. In contrast, more complex behaviour, such as grooming, required more standardisation of the illumination and video recordings to provide reliable results. Besides, the assessment of sucrose preference by measuring sucrose intake was particularly challenging. Sucrose solution contained blue dye to enable the visual inspection of the capillary in the video recordings - which was indistinguishable from the background when filled with a clear solution - and estimate the intake of sucrose by examining the abdomen of flies after behavioural testing.^25^ Although the blue dye made the capillary visible in the video recordings, it was useless to measure sucrose intake in most of the flies examined. In flies, the average consumption of fluids in 24 h may be in a range of 1 microliter (CAFE), ^22,23^ precluding reliable measurements of sucrose volume during the 5-min lane-maze test. Hence, the amount of sucrose intake by flies during the lane-maze test was uncertain in the present experiments. Therefore, sucrose preference was appraised by calculating an index of preference based on the time spent by flies on the capillary.

An index of preference was calculated considering the time of all activities that flies performed on the capillary, including locomotion, immobility, and grooming. The preference index could then be calculated using tracking data as obtained in ARC. ^27^ Raw measures of duration delivered much larger values of time in the lane than in the capillary because the capillary is smaller than the remainder of the lane, impeding a fair appraisal of the occupancy in different lane regions. Since the capillary occupies roughly 10% of a lane, it was expected that flies would spend around 10% of the time on the capillary and 90% in the remaining lane. Thus, the normalisation of the time spent by a fly in a lane sector by the length of the same sector (lane or capillary) is a more unbiased indicator of proportional exploration than raw time measurement. When normalised values were analysed, locomotion on the capillary was proportionally more frequent than on the remainder of the lane, while immobility and grooming were less prevalent and more equally distributed over the entire lane.

The difference between the expected and observed occupancy of the arena provided a measure for an individual or a group of flies *per se* without depending on an external reference, such as a control group. Positive, null, or negative values of the preference index may be interpreted as: preference, indifference (neutral), or avoidance of the capillary. Flies of all experimental groups spent proportionally more time near the capillary than in the other regions of the lane, indicating an attractive quality in the capillary region. Like in rodents ^14, 35, 36, 37, 38, 39, 40^, a combination of the different opportunities offered by the capillary region such as shelter, novelty and sucrose may contribute to the proportionally higher exploration of the capillary by flies in all groups. Future studies, e.g., “two-bottles” sucrose preference test, ^16^ should be performed to separate the preference for sucrose from other attractive features of the capillary.

According to previous literature, food deprivation may induce hyperactivity, *centrophilic* behaviour and sucrose preference in flies.^20,21^ In the present study, except for the small effect of food-deprivation for 8 h on the normalised times of grooming of females or immobility of males in the lane, no other obvious effect of food deprivation was observed. Indeed, food deprivation for 2, 8 or 20 h failed to promote hyperactivity compared to the control flies, maybe because locomotion is already a high incident behaviour in the lane-maze. Alternatively, the short duration of the test may explain the small effect sizes of food-deprivation on the motor activity of flies. The preference index revealed that the preference of flies for the capillary changed according to sex or experimental condition. The capillary preference seems neutral for male flies in most situations, except for the significantly high preference under control conditions (experiment 2) or food deprivation for two hours (experiment 1). In contrast, capillary preference seems neutral for female flies in most groups, except for the significantly high index in females deprived for 20 h.

In conclusion, data indicate that independent of sex or fasting conditions, flies explored more the capillary rather than the remainder of the lane. Food deprivation increased capillary preference in a sex or cohort-dependent fashion, which in females was more consistent than in males. Moreover, the duration of food deprivation in flies seems a determinant aspect modulating the preference for the capillary region of the lane. Data suggest that short lane-maze test is a feasibly high throughput assessment of sucrose preference in *D. melanogaster*, which may be sexually dimorphic as in other species studied so far.

Further studies are needed to confirm present findings overcoming limitations of the current study: 1-lack of power analysis, 2-short period of behavioural test, 3-use of anaesthesia to transfer flies to the lane-maze, 4-an indirect measure of the interaction of the fly with sucrose solution. Therefore, future studies should use power analysis to calculate sample sizes to detect the effects of food deprivation, or other intervention, on the preference index calculated from data scored in the lane-maze test. These future experiments should consider longer periods of the lane-maze test. To avoid the influence of the anaesthesia, flies may be transferred from the experimental tubes to lane-maze using a mouth aspirator as in other studies. ^31^ A practical solution to detect the interaction of the flies with the liquid sucrose would be to adapt a sensor for proboscis’s extension in the capillary. ^21^

## Acknowledgements

The authors thank Fabiani F Triches for the support in the maintenance of the vivarium, Tamires M Marchetto for the blinding of the video recordings of behavioural testing and João A Marcolan for the assistance with the Ethowatcher software. This manuscript was edited for English language by Ann C. Ferry.

## Declaration of conflicting interests

The authors declare that there is no conflict of interest.

## Funding

Fabiola B Eckert received fellowship from Conselho Nacional de Desenvolvimento Científico e Tecnológico, Brazil (140007/2016-4). This study was financed in part by the Coordenação de Aperfeiçoamento de Pessoal de Nível Superior – Brasil (CAPES) – Finance Code 001”.

## Notes

### Competing Interest Statement

The authors have declared no competing interest.

