## Supplementary material for "Lane-maze for preference testing in flies"

### Supplementary Methods:

#### Food for the flies: recipes for ten bottles of 300 ml

**Ingredients:** cornflour (58.34 g), sugar (45.84 g), soy flour (2.92 g), salt (0.45 g), rye flour (7.5 g) and antifungal agent (Nipagin) (1.6g) dissolved in 417 mL of filtered water.

**Procedures:** Boil the water, then add sugar, salt, antifungal agent and bring water to mild temperature. After that, add small portions of a mix of the flours to the water, mixing continuously and gently. Fill the bottles—cool down before transferring the flies to the medium. Alternatively, keep bottles with the medium in the refrigerator for up to 15 days.

#### Lane maze apparatus: recipe to print 3D

**Ingredients:** Acrylonitrile butadiene styrene filaments

**Procedures:** We used the OpenSCAD software to create the 3D project for the lane-maze for a 3D printer (GTMax3D Pro Core H4). After that, we used the slicer *Slic3r* in the Repetier-Host software to perform the slicing and generate the *gcode* file to print the maze in the 3D printer. Detailed information may be found at the Drosophila Information

Service [1] or in the OSF register of the publication [2] in the following link: https://osf.io/y46e9/files/osfstorage/5f6e0b0f9e9a3d02736e2d9a/

**Sample sizes, power analysis.**

No sample size calculation or power analysis was performed due to the lack of knowledge about suitable lane maze outcomes. The sample sizes were then selected based on studies performed with rodents, typically performed with sample sizes ranging from 6 to 20 individuals *per* group depending on the experimental designs,[3] and in preliminary studies in our laboratory.[4] Retrospective calculations of statistical power indicated that present studies were powered to detect large effect sizes (for calculations, we used the following link: https://www.ai-therapy.com/psychology-statistics/power-calculator).

**Detailed behavioural recording**

The ethographic analysis was performed using an open-source software package (Ethowatcher BetaOS 48)[5] developed in the laboratory of Bioengineering (CTC-IEB-UFSC) available upon request. The functionalities of the Ethowatcher BetaOS 48[5] are similar to the Ethowatcher [6] (freely available at https://ethowatcher.paginas.ufsc.br). A behavioural catalogue specifically created for the lane-maze test was used. The first version of the ethogram had items combining behavioural (locomotion, immobility, the motion of wings or legs) and spatial attributes of the lane (capillary region or lane subdivided in wall, floor, and roof). This catalogue was used in the first analysis, which results could be seen in Figure 2 of the manuscript and the description of the concordance analysis in Figure S1. According to the first analysis (locomotion, immobility, and grooming on the capillary region or in any place of the lane), the second version of the catalogue contained only the items with high reliability according to the first analysis (locomotion, immobility, and grooming on the capillary region or in any place of the lane). For each fly, Duration, latency, and frequency of each behaviour were extracted and pooled into periods of 5 minutes. In the initial analysis, behaviours were recorded from the entire content of the video recording of the test (40 min), while the final analysis was performed in five minutes of the test (between minutes $5^{th}$ and $10^{th}$). Transcriptions of behavioural categories from the video files were performed by a trained observer (FBE) and were carried out daily between 8 to 12 am, never exceeding four hours of continuous work. The video files were renamed by another person (JMN or TMM) to assure a blind assessment of the outcomes by the observer.  Concerning the behavioural recordings in the lane-maze test, the first step was to evaluate the quality and reliability of the assessment of each item of the catalogue using Cohen's *kappa* indexes, which may vary from almost perfect (upper 80%) to less than chance (0%)[7](Figure S1). The process for selection of the behavioural items comprised three steps: 1- qualitative appraisal to list all behaviours flies displayed in the lane-maze; 2-quantitative appraisal (performed twice) to register the latency (s), frequency, and duration (s) of all behaviours listed in the first step, analysing nine videos randomly selected; 3-calculation of Cohen's *kappa* index for each item using the automatic function of the Ethowatcher BetaOS 48.

**Figure S1:** Agreement calculations

a) $$K = \frac{P_O - P_E}{1 - P_E}$$

b) $$P_O = \frac{\text{Agreements}}{\text{Agreements + Disagreements}}$$

c) $$P_E = \sum_{J=1}^{n} \frac{P_{1k} \times P_{2k}}{\text{Agreements + Disagreements}}$$

**Legend for Figure S1:** Agreement calculations. a) Cohen's *kappa* index. b) Proportion of total agreements between observers. c) Proportion of agreements by chance. Abbreviations: 1k= kappa index; 2k= kappa index; E= expected; O= observed; P= proportion.

**Calculation of normalised duration:**

Normalised durations (Nd) consisted of dividing the time (s) spent in either sector (lane or capillary) by the length (mm) of the respective sector of the lane (lane or capillary) (Figure S2). The total length of the lane, including the capillary area, is 50 mm. The capillary area (mm) length was variable from lane to lane and from test to test due to variations in the insertion procedures. Lengths (mm) of the capillaries were extracted from video recordings with a digital ruler of the Ethowatcher BetaOS 48.

**Figure S2:** Normalised durations calculation

a)

$$Nd = \frac{\text{duration (s) in the lane}}{\text{(50 mm - length of the capillary (mm))}}$$

b)

$$Nd = \frac{\text{duration (s) in the capillary}}{\text{length of the capillary (mm)}}$$

**Legend for Figure S2:** Normalised durations calculations. a) Calculation of normalised durations in the lane. b) Calculation of normalised durations in the capillary. Abbreviations: Nd= normalised duration.

**Calculation of preference index:**

Preference index ($P_{index}$) consisted of estimate the "expected" time flies spent in a sector of the lane (lane or capillary) in the absence of preference (Figure S3 a and b) and then subtract it from the "observed" time (Nd, Figure S2) (Figure S3c). $P_{index}$ may be positive, or null or negative values indicating higher, similar, or lower occupancy of the capillary region. The occupancy of the capillary region was interpreted as "preference" to the capillary region. The higher the occupancy, the higher the preference for the capillary. Therefore, positive values of $P_{index}$ indicate "preference" while "negative" values indicate aversion for the capillary. The null value of the $P_{index}$ indicates an absence of preference, i.e., the capillary is neutral. $P_{index}$

was expressed as a percentage ($P_{index\ (\%)}$, Figure S3 d) to take the duration of behavioural testing into account.

**Figure S3:** Calculation of preference index

a) $$\text{Proportion (mm)} = \frac{(\text{length of the capillary (mm)} \times 100)}{50\text{mm}}$$

b) $$\text{Expected time (E)} = \frac{\text{proportion (mm)} \times 300\text{s}}{100\%}$$

c) $$P_{index} = \text{Observed time (O)} - \text{Expected time (E)}$$

d) $$P_{index\ (\%)} = \frac{(\text{O} - \text{E}) \times 100}{300\text{s}}$$

**Legend for Figure S3:** Calculation of preference index. a) Calculation of proportion occupancy of the capillary in the lane. b) Calculation of expected time (s) in the capillary. c) Calculation of preference index based on the observed and expected time (s) in the capillary. d) Transformation of preference index in percentage. Abbreviations: $P_{index}$ = preference index; E= expected time; O=observed time.

**Supplementary Results:**

**Behavioural catalogue, quality, and reliability of behavioural assessment:**

The behavioural catalogue used for assessing behavioural outcomes in experiments 1-3 was created based on the quality and reliability of behavioural assessments (Cohen’s *kappa* index, calculations in Figure S1). In the experiments, 1-3, only those behavioural items with a substantial agreement between the measures (Cohen's *kappa* index upper 75%) were included in the behavioural catalogue. In general, flies walked (locomotion) or stayed in a place completely immobile (immobility) or moving body parts (legs or wings) in any place of the lane (floor, wall, roof, capillary). The motion of the wings, especially on the capillary, was a rare event. Thus, the "motion of wings" and "motion of legs" were merged into the single item named "grooming" for the subsequent analysis. Reliability of the items registered varied from item to item, as follows (decreasing order of Cohen’s *kappa* indexes, Figure S4): immobility on the floor (92±3%), locomotion on the wall (90 ±0.55%); locomotion on capillary (86±1%); motion of legs on the wall (78±4%); motion of wings on the wall (75±5%), immobility on the wall (74±7%); locomotion on the roof (74 ±4%); immobility on the roof (73±10%); immobility on the capillary (68±10%); locomotion on the floor (44±16%). A second analysis of concordance, suppressing specific attributes of the lane (floor, wall, or roof), increased the reliability for all items of the catalogue to values upper 80% (almost perfect)[7]. Therefore, behavioural assessments in experiments 1-3 were performed with a behavioural catalogue containing the following items (description in Table S2): immobility in the lane; locomotion in the lane; grooming in the lane; immobility on the capillary; locomotion on the capillary; grooming on the capillary.

**Figure S4:** Intra-observer agreement and agreement by behaviour

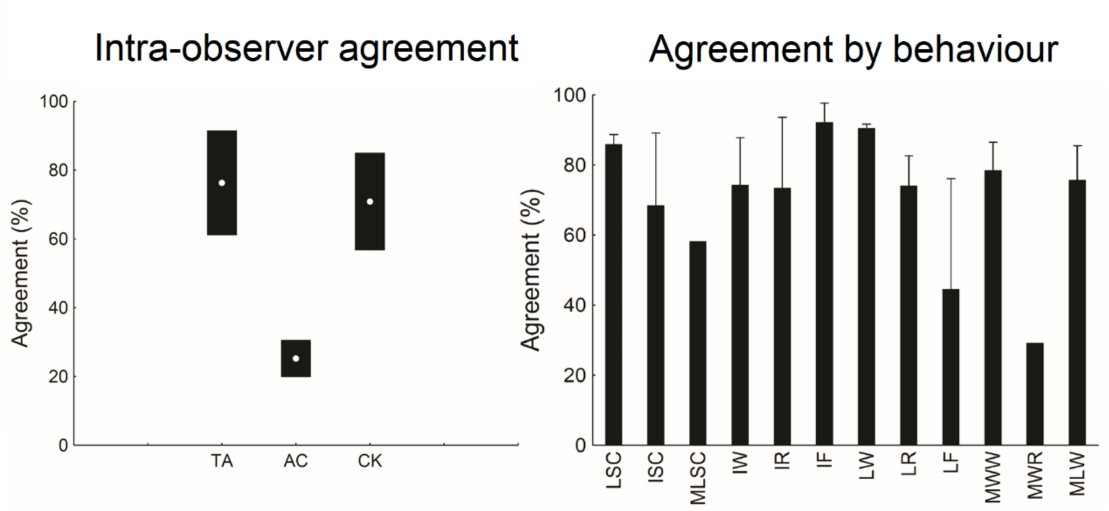


**Legend for Figure S4:** Intra-observer agreement (left panel) and agreement by behaviour (right panel). Data are expressed as mean ±SEM. Abbreviations: TA (total frequency-based % agreement), AC (agreement by chance), CK (Cohen's *kappa*), LSC(locomotion on sucrose capillary), ISC (immobility on sucrose capillary), MLSC (move legs on sucrose capillary), IW (immobility on the wall), IF (immobility on the floor), LW (locomotion on the wall), LR(locomotion on the roof), LF (locomotion on the floor), MWW(move wings on the wall), MWR (move wings on the roof) and MLW (move legs on the wall).

Table S1: Agreement by behaviour (%) – raw data

| Category | sample size | Video recordings displaying a given category | Mean (% agreement by behaviour) | SD |
|---|---|---|---|---|
| LSC | 9 | 9 | 85.97726 | 4.104893 |
| ISC | 9 | 4 | 68.44285 | 21.09251 |
| MLSC | 9 | 1 | 58.17490 | - |
| IW | 9 | 9 | 74.33189 | 20.55013 |
| IR | 9 | 6 | 73.42410 | 25.19232 |
| IF | 9 | 8 | 92.18130 | 7.888412 |
| LW | 9 | 9 | 90.53671 | 1.670068 |
| LR | 9 | 8 | 74.02559 | 12.35068 |
| LF | 9 | 5 | 44.59038 | 35.92329 |
| MWW | 9 | 9 | 78.43130 | 12.32817 |
| MWR | 9 | 1 | 29.25532 | - |
| MLW | 9 | 9 | 75.69590 | 15.03803 |

**Legend for Table S1:** Data are expressed as mean ±SD. Abbreviations: LSC (locomotion on sucrose capillary), ISC (immobility on sucrose capillary), MLSC (move legs on sucrose capillary), IW (immobility on the wall), IF (immobility on the floor), LW (locomotion on the wall), LR (locomotion on the roof), LF (locomotion on the floor), MWW (move wings on the wall), MWR (move wings on the roof) and MLW (move legs on the wall).

Table S2: Final version of the behavioural catalogue

| Location | Behaviour | Behaviour description |
|---|---|---|
| Capillary | Locomotion | The fly moves its entire body using leg movements in frontal, lateral or backward progression above, beside or in front of the capillary containing sucrose. |
| | Immobility | The fly stays still, without moving any part of the body that can be observed and not move from the place where it is, staying on top, beside or in front of the capillary containing sucrose. |
| | Grooming | The fly does not move its body from where it is, only moves the wings or legs above, beside or in front of the capillary containing sucrose. |
| Lane | Locomotion | The fly moves its entire body using leg movements in frontal, lateral or backward progression above on the lane. |
| | Immobility | Fly stays still, without moving any part of the body that can be observed and not move from the place where it is, on the lane. |
| | Grooming | Fly does not move its body from where it is and only moves the wings or legs, on the lane. |

**Analysis of the pilot studies:**

Preliminary analyses were performed using data of flies from control groups. After each behavioural test in the lane-maze, flies were examined under a stereoscopic microscope to detect the presence of blue dye in their abdomens. Due to insufficient microscopic power or the low concentration of blue dye in the abdomens of the flies, the poor performance of this method precludes further analysis of these data.

Duration (s) of behaviours was arbitrarily selected to analyse experiments 1-3. Latencies and frequencies were not analysed. The analysis of the whole duration of the test (40 min) indicated that locomotion duration remained stable over the test while grooming decreased, and immobility increased over time. Grooming duration decreased steadily from 15$^{th}$ min reaching until the end of the test. Immobility duration increased steadily from 15$^{th}$ min, reaching a plateau until the end of the test. Therefore, the time-window between the 5$^{th}$ and 10$^{th}$ min of the lane-maze test was selected to allow for effects of the factors such as sex or fasting conditions on locomotion, grooming or immobility. See Table S4 for means, standard errors or standard deviations for every experimental group in the independent experiment. These raw data were normalised to represent the time for every behavioural outcome registered in the lane or capillary proportionally (Figure 3, in the manuscript). Most raw or normalised data failed to present normal distribution and homogeneity of variance, even with different transformations, demanding adopting a non-parametric statistical analysis to compare data from control and experimental groups in the independent experiments.

**Table S3: Duration (s) of locomotion, immobility and grooming in the pilot study.**

| | | Capillary | | | Lane | | |
|---|---|---|---|---|---|---|---|
| Sex | Period (min) | Locomotion | Immobility | Grooming | Locomotion | Immobility | Grooming |
| Female control (n=30) | 5 | 17.0±19.2 | 7.9±38.6 | 1.1±4.8 | 85.3±64.5 | 154.1±81.7 | 33.6±48.3 |
| | 10 | 22.0±14.2 | 1.4±3.6 | 1.2±6.7 | 138.4±64.8 | 44.1±44.5 | 92.8±66.0 |
| | 15 | 19.0±16.6 | 1.7±4.1 | 2.3±12.0 | 108.8±67.4 | 44.9±43.5 | 123.5±73.2 |
| | 20 | 19.0±19.0 | 4.7±10.6 | 2.3±11.1 | 100.0±66.2 | 72.8±61.1 | 101.0±70.0 |
| | 25 | 21.2±18.5 | 0.8±3.9 | 0.0±0.0 | 108.2±74.2 | 98.3±80.2 | 71.3±63.8 |
| | 30 | 23.0±20.0 | 0.8±2.2 | 0.7±3.3 | 110.5±71.0 | 111.7±91.5 | 53.3±52.7 |
| | 35 | 21.1±18.3 | 0.4±1.2 | 7.1±40.0 | 106.9±71.8 | 111.4±86.9 | 53.0±62.0 |
| | 40 | 21.0±20.1 | 2.5±7.2 | 1.7±9.1 | 91.5±69.0 | 117.0±93.5 | 30.1±47.5 |
| Male control (n=28) | 5 | 13.4±14.5 | 2.2±6.7 | 0.4±2.0 | 91.5±57.1 | 162.8±65.4 | 28.6±27.4 |
| | 10 | 18.9±14.3 | 3.3±10.4 | 0.3±1.6 | 120.6±64.7 | 61.4±63.8 | 95.4±63.8 |
| | 15 | 15.7±12.3 | 0.3±1.3 | 0.6±3.4 | 98.1±61.2 | 63.0±67.2 | 121.1±56.8 |
| | 20 | 14.8±11.8 | 5.2±22.6 | 1.1±5.8 | 92.0±58.4 | 101.7±76.5 | 85.0±52.1 |
| | 25 | 16.4±12.5 | 0.8±2.2 | 0.0±0.0 | 94.6±59.7 | 119.0±87.0 | 69.0±64.6 |
| | 30 | 16.6±14.5 | 5.3±25.7 | 1.8±7.0 | 89.8±65.7 | 139.1±95.3 | 47.2±73.6 |
| | 35 | 23.8±16.3 | 16.4±48.7 | 0.0±0.0 | 117.8±66.3 | 110.6±75.0 | 31.4±37.1 |
| | 40 | 24.5±14.9 | 16.3±43.3 | 0.0±0.0 | 115.0±64.7 | 109.8±80.2 | 26.3±42.3 |

Legend for Table S3: Duration (s) of locomotion, immobility and grooming in the lane or on the capillary of females (n=30) or males (n=28) flies kept in standard laboratory conditions before the lane-maze test (40 min). Means±SD *per period* (raw data). Abbreviations: min= minute; n= sample size.

**Statistics for Experiment 1:**

No significant differences were observed among the groups (male or female flies in standard conditions or food deprivation 2 h, Figure S5) when analysing either raw or normalised durations of locomotion, immobility or grooming that were assessed during the 5-min of the lane-maze test. For the raw durations on the capillary, no significant differences were seen among the groups for locomotion ($H(3) = 3.18$, $p = .36$), immobility ($H(3) = 3.68$, $p = .30$) or grooming ($H(3) = 2.35$, $p = .50$). In the lane, no significant differences were seen among the groups for the raw duration of locomotion ($H(3) = 4.85$, $p = .18$), immobility ($H(3) = 1.10$, $p = .77$), grooming ($H(3) = 4.24$, $p = .23$). Normalised durations were not conclusively different among the groups for any category in the lane (locomotion: $H(3) = 5.59$, $p = .13$; immobility: $H(3) = 1.10$, $p = .78$; grooming: $H(3) = 4.12$, $p = .25$) or capillary (locomotion: $H(3) = 2.42$, $p = .49$; immobility: $H(3) = 4.19$, $p = .24$; grooming: $H(3) = 2.35$, $p = .50$) (Figure S5).

**Figure S5:** Duration (s) of locomotion, immobility and grooming in experiment 1.

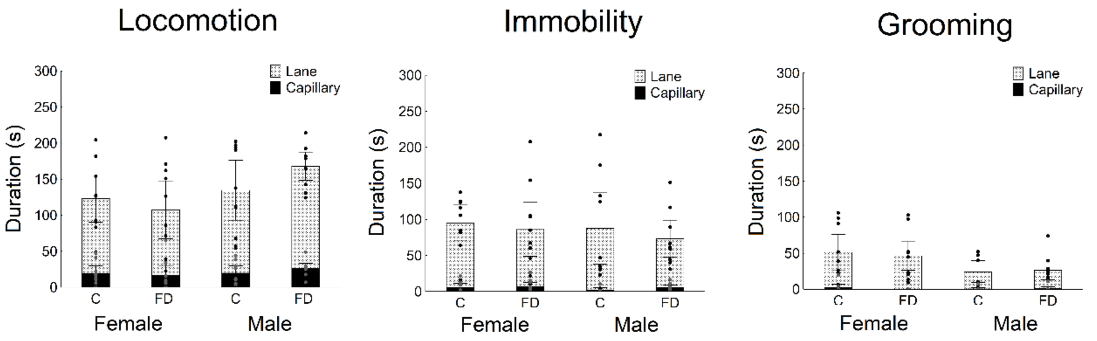


**Legend for Figure S5:** Duration of locomotion (left panels), immobility (middle panels), grooming (right panels) of female or male flies of the control (females, n=9; males, n=9) or food-deprived for 2 h (females, n=10; males, n=9) groups on the capillary (black bars) or lane (grey bars) during 5 min of the lane-maze test. Data are expressed as mean ±SEM. Abbreviations: FD= food deprivation, C=control, F= female, M=male. Grey circles= raw data on the capillary, black circles= raw data on the lane.

**Statistics for Experiment 2:**

For raw durations on the capillary, no significant differences were seen among the groups for locomotion (H (3) =2.97, *p* =.39), immobility (H (3) =4.025, *p* =.26) and grooming (H (3) =1.11, *p* =.77). In the lane, no significant differences were seen among the groups for the raw duration of locomotion (H (3) =1.42, *p* =.70), immobility (H (3) =7.70, *p* =.052) and grooming (H (3) =4.92, *p* = .17). Normalised durations were not conclusively different among most of the groups for any category on capillary (locomotion: H (3) =4.13, $p$ = .24; immobility: H (3) =4.46, $p$ = .21; grooming: H (3) =1.11, $p$ = .77) or locomotion in the lane (H (3) =1.37, $p$ = .71) (Figure S6).

**Figure S6:** Duration (s) of locomotion, immobility and grooming in experiment 2.

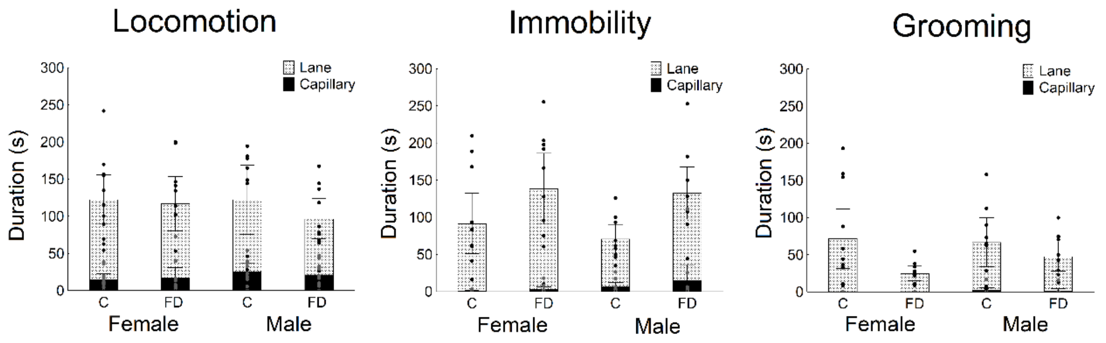


**Legend for Figure S6:** Duration of locomotion (left panels), immobility (middle panels), grooming (right panels) of female or male flies of the control (females, n=11; males, n=9) or food-deprived for 8 h (females, n=10; males, n=10) groups on the capillary (black bars) or lane (grey bars) during 5 min of the lane-maze test. Data are expressed as mean ±SEM. Abbreviations: FD= food deprivation, C=control, F= female, M=male. Grey circles= raw data on the capillary, black circles= raw data on the lane

**Statistics for Experiment 3:**

No significant differences were observed among the groups (male or female flies in standard conditions or food deprivation 20 h, Figure S7) when analysing either raw or normalised durations assessed during the 5$^{th}$ and 10$^{th}$ min of the lane-maze test. For raw durations on the capillary, no significant differences were seen among the groups for locomotion (H (3) =6.41, $p$ = .09), immobility (H (3) =5.93, $p$ = .11) and grooming (H (3) =2.05, $p$ = .56). In the lane, no significant differences were seen among the groups for raw durations of locomotion (H (3) =5.61, $p$ = .13), immobility (H (3) =4.25, $p$ = .23) and grooming (H (3) =2.89, $p$ = .41). Normalised durations were not conclusively different among the groups for any category in the lane (locomotion: H (3) =5.74, $p$ = .12; immobility: H (3) =4.47, $p$ = .21; grooming: H (3) =5.56, $p$ = .13) or capillary (locomotion: H (3) =6.39, $p$ = .09; immobility: H (3) =5.59, $p$ = .13; grooming: H (3) =2.05, $p$ = .56) (Figure S7).

**Figure S7:** Duration (s) of locomotion, immobility and grooming in experiment 3.

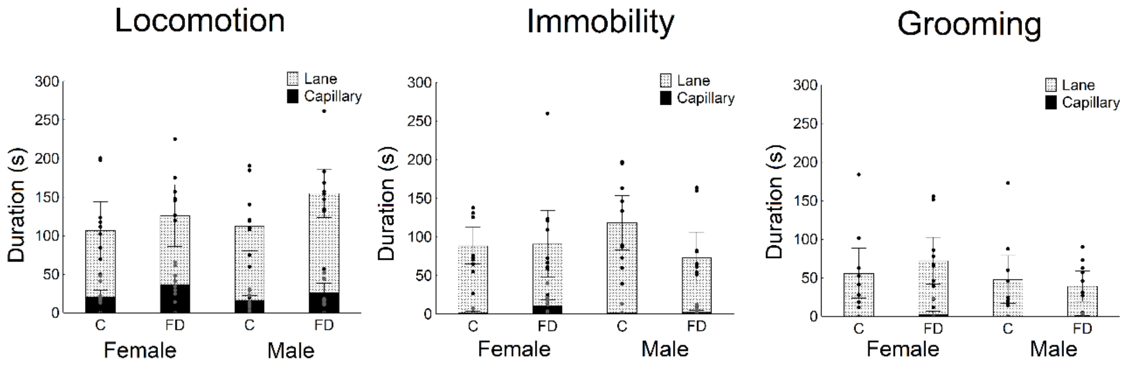


**Legend for Figure S7:** Duration of locomotion (left panels), immobility (middle panels), grooming (right panels) of female or male flies of the control (females, n=10; males, n=10) or food-deprived for 20h (females, n=10; males, n=10) groups on the capillary (black bars) or lane (grey bars) during 5 min of the lane-maze test. Data are expressed as mean ±SEM. Abbreviations: FD= food deprivation, C=control, F= female, M=male. Grey circles= raw data on the capillary, black circles= raw data on the lane.

**Data with standard deviations**

**Table S4:** Duration (s) of behaviours recorded in the first 5 min of the lane-maze test in experiments 1, 2 and 3

| Sex | Group (n) | Capillary | | | | | | | | | Lane | | | | | | | | |
|---|---|---|---|---|---|---|---|---|---|---|---|---|---|---|---|---|---|---|---|
| | | Locomotion | | | Immobility | | | Grooming | | | Locomotion | | | Immobility | | | Grooming | | |
| | | Mean | SEM | SD | Mean | SEM | SD | Mean | SEM | SD | Mean | SEM | SD | Mean | SEM | SD | Mean | SEM | SD |
| F | C 2h (n=9) | 19.4 | 5.2 | 15.5 | 6.3 | 2.5 | 7.5 | 2.6 | 2.2* | 6.7 | 123.3 | 16.7 | 50.0 | 95.0 | 12.6 | 38.0 | 51.5 | 12.5 | 37.7 |
| M | C 2h (n=9) | 20.0 | 5.0 | 14.8 | 2.4 | 1.5 | 4.5 | 0.8 | 0.8 | 2.4 | 134.7 | 21.2* | 63.8 | 87.8 | 25.5* | 76.4 | 24.4 | 7.6 | 23.0 |
| F | FD 2h (n=10) | 17.0 | 6.6 | 21.0 | 7.2 | 2.8 | 8.7 | 0.0 | 0.0* | 0.0 | 107.3 | 20.5 | 65.0 | 86.3 | 19.0 | 60.1 | 46.6 | 10.3 | 32.5 |
| M | FD2h (n=9) | 26.4 | 3.7 | 11.1 | 5.5 | 1.8 | 5.4 | 1.2 | 1.2 | 3.6 | 167.9 | 10.0* | 30.0 | 73.3 | 13.1* | 39.3 | 26.7 | 6.9 | 20.7 |
| F | C 8h (n=11) | 15.2 | 3.6 | 12.1 | 0.8 | 0.4 | 1.4 | 0.0 | 0.0* | 0.0 | 122.6 | 17.0 | 56.1 | 91.6 | 20.8 | 69.0 | 71.4 | 20.4* | 67.9 |
| M | C 8h (n=9) | 25.0 | 5.8 | 17.3 | 6.4 | 3.1 | 9.4 | 1.9 | 1.9 | 5.7 | 122.4 | 23.7* | 71.1 | 71.0 | 10.0 | 29.6 | 66.7 | 16.6 | 50.0 |
| F | FD 8h (n=10) | 17.0 | 7.1 | 22.5 | 3.0 | 1.9 | 5.9 | 0.0 | 0.0* | 0.0 | 117.0 | 18.7 | 59.1 | 138.4 | 24.4 | 77.1 | 24.8 | 4.9* | 15.8 |
| M | FD 8h (n=10) | 21.2 | 5.1 | 16.3 | 14.8 | 11.0 | 34.7 | 1.6 | 16.0 | 5.0 | 96.4 | 13.8* | 43.6 | 132.7 | 17.8 | 56.4 | 47.6 | 10.0 | 31.9 |
| F | C 20h (n=10) | 21.1 | 4.4 | 13.8 | 1.5 | 1.0* | 3.1 | 0.0 | 0.0* | 0.0 | 106.6 | 19.0 | 59.8 | 88.4 | 12.1 | 38.3 | 55.9 | 16.5 | 52.1 |
| M | C 20h (n=10) | 16.0 | 3.5 | 11.1 | 1.6 | 1.3* | 4.2 | 0.0 | 0.0 | 0.0 | 112.7 | 16.6 | 52.7 | 118.3 | 17.9 | 56.5 | 47.9 | 15.8* | 50.0 |
| F | FD 20h (n=10) | 36.6 | 6.5 | 20.4 | 10.7 | 4.1* | 13.0 | 2.3 | 2.3* | 7.3 | 125.8 | 20.4 | 64.6 | 90.9 | 22.2 | 70.1 | 72.3 | 15.4 | 48.7 |
| M | FD 20h (n=10) | 26.6 | 6.1 | 19.2 | 2.4 | 1.3* | 4.2 | 0.0 | 0.0 | 0.0 | 155.0 | 16.0 | 50.5 | 72.8 | 17.0 | 53.9 | 39.3 | 10.0* | 31.6 |

**Legend for Table S4:** Means *per group* (raw data). Abbreviations: SEM= standard error of mean; SD= standard deviation of mean; C= control group; F= females; FD= food deprivation group; h= hours; M=male; n= sample size. Flies of FD groups were food-deprived for two h or eight h or 20 h. *= Levene test, p <0.05.

Table S5: Proportion (seconds *per* millimetres, s/mm) of each behaviour.

| | | Capillary | | | Lane | | |
|---|---|---|---|---|---|---|---|
| Sex | Group (n) | Locomotion | Immobility | Grooming | Locomotion | Immobility | Grooming |
| F | C 2h (n=9) | 6.0±4.5 | 2.3±3.5 | 0.8±2.2 | 2.6±1.0 | 2.0±0.8 | 1.1±0.8 |
| M | C 2h (n=9) | 5.2±3.6 | 0.5±.0.9 | 0.2±0.6 | 2.9±1.4 | 1.9±1.6 | 1.1±0.6 |
| F | FD 2h (n=10) | 4.7±5.1 | 2.1±2.1 | 0.0±0.0 | 2.3±1.4 | 1.8±1.3 | 1.5±1.5 |
| M | FD2h (n=9) | 7.1±2.4 | 1.4±1.3 | 0.2±0.7 | 3.6±0.6 | 1.6±0.8 | 0.5±0.3 |
| F | C 8h (n=11) | 4.4±3.6 | 0.2±0.4 | 0.0±0.0 | 2.6±1.2 | 1.9±1.5 | 1.2±1.0 |
| M | C 8h (n=9) | 8.3±5.8 | 2.1±3.1 | 0.6±1.9 | 2.6±1.5 | 1.5±0.6 | 1.5±1.0 |
| F | FD 8h (n=10) | 4.4±5.5 | 0.7±1.5 | 0.0±0.0 | 2.5±1.2 | 3.0±1.6 | 0.5±0.5 |
| M | FD 8h (n=10) | 6.1±4.6 | 3.8±8.6 | 0.4±1.2 | 2.0±0.9 | 2.8±1.2 | 0.7±0.5 |
| F | C 20h (n=10) | 6.3±4.0 | 0.3±0.8 | 0.0±0.0 | 2.3±1.3 | 1.9±0.8 | 1.7±1.2 |
| M | C 20h (n=10) | 5.1±3.4 | 0.5±1.4 | 0.0±0.0 | 2.4±1.1 | 2.5±1.2 | 1.0±0.6 |
| F | FD 20h (n=10) | 11.1±6.1 | 3.1±4.1 | 0.4±1.4 | 2.7±1.4 | 1.9±1.5 | 0.7±0.5 |
| M | FD 20h (n=10) | 9.8±8.0 | 0.9±1.4 | 0.1±0.4 | 3.3±1.0 | 1.5±1.1 | 0.8±0.6 |

Legend for Table S5: Proportion (seconds *per* millimetres, s/mm) of locomotion, immobility, and grooming of female or male flies of the control (C) or food-deprived (FD) groups on the capillary or lane during 5 min of the lane-maze test. Data are expressed as mean ± SD. Abbreviations: FD= food deprivation, C=control, F= female, M=male; h= hours; n= sample size.

Table S6: Preference for the capillary region of the lane index (%)

| Sex | Group (n) | Preference index (%) |
|---|---|---|
| F | C 2h (n=9) | 2.5±7.2 |
| M | C 2h (n=9) | 0.2±4.6 |
| F | FD 2h (n=10) | 1.4±8.1 |
| M | FD2h (n=9) | 3.7±5.4 |
| F | C 8h (n=11) | -1.5±3.9 |
| M | C 8h (n=9) | 5.1±6.8 |
| F | FD 8h (n=10) | -0.5±8.8 |
| M | FD 8h (n=10) | 5.7±14.6 |
| F | C 20h (n=10) | 0.7±5.1 |
| M | C 20h (n=10) | -0.1±3.8 |
| F | FD 20h (n=10) | 9.9±9.5 |
| M | FD 20h (n=10) | 4.0±7.0 |

Legend for Table S6: Preference for the capillary region of the lane index (%) of 2h groups, 8h groups and 20h groups. Data are expressed as mean ±SEM. Abbreviations: FD= food deprivation, C=control, F= female, M=male; h=hours, n= sample size. Data are expressed as mean ± SD.
